# Rice Galaxy: an open resource for plant science

**DOI:** 10.1101/358754

**Authors:** Venice Juanillas, Alexis Dereeper, Nicolas Beaume, Gaetan Droc, Joshua Dizon, John Robert Mendoza, Jon Peter Perdon, Locedie Mansueto, Lindsay Triplett, Jillian Lang, Gabriel Zhou, Kunalan Ratharanjan, Beth Plale, Jason Haga, Jan E. Leach, Manuel Ruiz, Michael Thomson, Nickolai Alexandrov, Pierre Larmande, Tobias Kretzschmar, Ramil P. Mauleon

## Abstract

*Background*

Rice molecular genetics, breeding, genetic diversity, and allied research (such as rice-pathogen interaction) have adopted sequencing technologies and high density genotyping platforms for genome variation analysis and gene discovery. Germplasm collections representing rice diversity, improved varieties and elite breeding materials are accessible through rice gene banks for use in research and breeding, with many having genome sequences and high density genotype data available. Combining phenotypic and genotypic information on these accessions enables genome-wide association analysis, which is driving quantitative trait loci (QTL) discovery and molecular marker development. Comparative sequence analyses across QTL regions facilitate the discovery of novel alleles. Analyses involving DNA sequences and large genotyping matrices for thousands of samples, however, pose a challenge to non-computer savvy rice researchers.

*Findings*

We adopted the Galaxy framework to build the federated Rice Galaxy resource, with shared datasets, tools, and analysis workflows relevant to rice research. The shared datasets include high density genotypes from the 3,000 Rice Genomes project and sequences with corresponding annotations from nine published rice genomes. Rice Galaxy includes tools for designing single nucleotide polymorphism (SNP) assays, analyzing genome-wide association studies, population diversity, rice-bacterial pathogen diagnostics, and a suite of published genomic prediction methods. A prototype Rice Galaxy compliant to Open Access, Open Data, and Findable, Accessible, Interoperable, and Reproducible principles is also presented.

*Conclusions*

Rice Galaxy is a freely available resource that empowers the plant research community to perform state-of-the-art analyses and utilize publicly available big datasets for both fundamental and applied science.

## Findings

### Background

With the decreasing cost of genome sequencing, rice molecular geneticists, breeders and diversity researchers are increasingly adopting genotyping technologies as routine components in their workflows, generating large datasets of genotyping and genome sequence information. Concurrently international consortia have made re-sequencing or high density genotyping data from representative diversity collections publically available. These include, but are not limited to the medium-depth (15-20x coverage) resequencing data of the 3,010 accessions from the 3K Rice Genome (3K RG) Project (˜1 – 2 million SNPs per accession) [1,2] and the 700,000 SNP Affymetrix array data for the 1,445 accessions of the High Density Rice Array (HDRA) germplasm collections [3]. The corresponding accessions are available at non-profit prices from the Genetic Resource Center (GRC) of the International Rice Research Institute (IRRI) for phenotyping, allowing subsequent Genome-Wide Association Studies (GWAS). Analysis of such datasets is a challenge to rice researchers due to (1) the fairly large data matrix and the compute-intensive algorithms that requires specialized computing infrastructure (a fairly large RAM, powerful CPU, and large disk space), and (2) the relative difficulty in using Open Source / free software tools for analysis, which are commonly provided without graphical user interface and require proper installation in a Linux operating system environment.

On the computational side, public web resources with specialized tools already exist, and are maintained at different institutions. The Rice SNP-Seek database [4,5], largely developed and hosted by IRRI, contains phenotypic, genotypic, and passport information for over 4,400 rice accessions from large scale rice diversity projects such as the 3K RG and the HDRA collections. SNP-Seek (http://snp-seek.irri.org) currently contains phenotype data for 70 different morphological and agronomic traits and stores SNPs and small indels discovered by mapping the 3K RG accessions to four published rice draft genome assemblies, collectively resulting in the discovery of ˜11M new SNPs and ˜0.5M new indels. While SNP-Seek focused on delivery of prior analyzed content rather than providing an analysis platform, Gigwa [6] (http://gigwa.southgreen.fr/gigwa/), hosted at the South Green portal [7] (http://www.southgreen.fr/), is a scalable and user-friendly web-based tool which provides an easy and intuitive way to explore large amounts of genotyping data from next-generation sequencing (NGS) experiments. Gigwa allows for filtering of genomic and genotyping data from NGS analyses based not only on variant features, including functional annotations, but also on genotype patterns to explore the structure of genomes in an evolutionary context for a better understanding of the ecological adaptation of organisms. Gramene [8] is a curated, open-source, integrated data resource for comparative functional genomics in crops and model plant species that, among other species, includes rice. Data and analysis tools are available as portals at the Gramene site (http://gramene.org/). In these resources mentioned, the analyses methodologies are custom-built by the respective projects.

There are other freely available web-based bioinformatics and breeding informatics software tools, optimized for plant species other than rice, including Araport (https://www.araport.org/) for Arabidopsis, Cassavabase (https://cassavabase.org/) for cassava, and The Triticeae Toolbox (T3, https://triticeaetoolbox.org/) for wheat and barley. While these tools are very useful, they are species/crop specific and custom-built for the specialized requirements of their respective communities (such as project datasets), making adoption in rice challenging for at least two reasons: (1) the need to produce curated rice datasets that work seamlessly with the software system (e.g. genome-browser ready data, curated genes, published QTLs from biparental crosses and GWAS and markers associated to traits), and (2) the need for a dedicated software development team to customize the application for rice-specific data and analyses.

The ability of software to automate repetitive analyses task is attractive for data analysts, and the public sharing of the analytical methodology (as opposed to just the raw data and the results) enhances reproducibility and is being supported by academic communities of practice such as FORCE11 (https://www.force11.org). Many research groups working with NGS data have a high demand for computing infrastructure and their complex analyses often comprise several steps using different software tools (pipeline). The deployment of these different software tools is a big challenge to small institutions without dedicated scientific computing support staff. There is no single solution to address these challenges. Our approach to help overcome them is the integration of a range of these different bioinformatics tools into the Galaxy bioinformatics system. Galaxy [9] is a web-based analysis workbench and workflow management system initiated at the Penn State University. It includes a collection of software packages which can be operated via a web browser on a public server. Galaxy is a mature community effort, supported by various high-powered institutions, is relatively easy to deploy and maintain, and thus well-suited to serve low and moderately resourced institutions such as IRRI. The graphical user interface of Galaxy means that no knowledge of code is needed, thus facilitating bioinformatics analyses by researches without computational expertise.

We built a suite of federated Galaxy resources and tools, which we collectively named **Rice Galaxy** (Figure 1). Rice Galaxy contains shared software tools and datasets tailored to the needs of rice researchers and breeders, providing computing resources through an easy-to-use interface, and allowing reproducibility and publication of analytical methodology and results.

**Figure 1.**
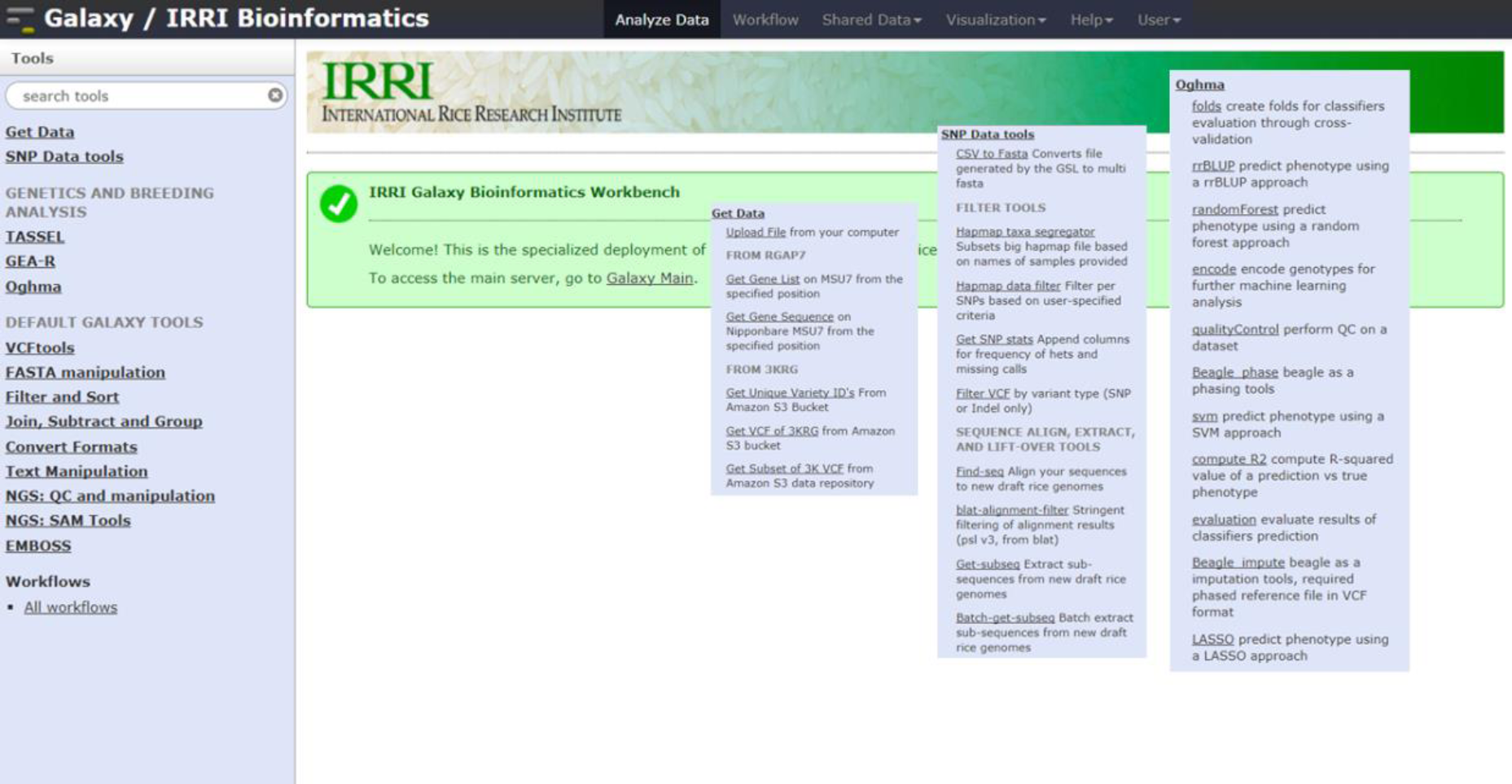
Rice Galaxy @ IRRI with customized analyses tools for genetics, breeding, and custom data sources (i.e. 3,000 Rice Genomes project).

The Rice Galaxy federated resources are hosted at:

- IRRI Galaxy @ International Rice Research Institute: http://galaxy.irri.org
- Rice Galaxy (common) Toolshed: http://52.76.88.51:8081/

## Discussion

### 1. Built-in / interoperable rice data

The Rice Galaxy system is customized to provide rice-specific genomic and genotypic data. Of primary importance is the gold-standard *japonica* variety reference genome (Nipponbare IRGSP release 1.0) [10], to which the reference gene models and most of the SNPs published have been anchored. In addition, eight medium to high quality published genomes from various sequencing projects and the respective genome annotations for each are installed as alternative genome builds and are available as drop-down menu choices in Rice Galaxy. These include four high-quality builds from *indica*-type varieties Minghui 63 and Zhengshan 97 [11], IR 8 (GenBank: MPPV00000000.1), Shuhui 498 [12], as well as an aus-type variety N 22 (GenBank: LWDA00000000.1), as well as four medium to low quality genomes, two *indica* (IR 64, [13] and 93-11, [14]) and two aus-type rice genomes (DJ 123, [13] and Kasalath, [15]). While these references were selected to represent diversity, they further represent variety groups that display agronomically important characteristics, such as heat and drought tolerance, disease resistance, submergence tolerance, adaptation to low-phosphorus soil, wide adaptability, good grain quality, aerobic (upland) adaptation and deep roots [16–18]. Even though these genomes are highly similar to each other, they each contain unique regions (from 12.3 Mbp to 79.6 Mbp) that may harbor genes restricted to these variety-groups [5]. With the availability of several reference genomes, it becomes relatively straightforward to custom design SNP assays that are either of broad utility across varietal groups or specific to single groups.

Rice Galaxy includes genotyping data of the 3k RG (such as the 3K RG 3024 accessions x 4.8M filtered SNPs, 440K core SNPs, 1M GWAS-ready SNPs, and 2.3M indels) useful for GWAS, region-specific diversity analyses, and single locus allele mining in the shared data library.

### 2. Toolkits Built (and detailed discussion of each toolkit)

#### SNP assay design: Lift-over of SNPs from one genome to another

SNPs discovered relative to the gold-standard reference genome (Nipponbare IRGSP 1.0, [10]) are commonly used in QTL mapping (either by GWAS or biparental cross). In order to develop robust markers associated with the trait of interest, however, a SNP assay that works in the target varietal groups is needed. Consequently there is a need to “lift-over” SNPs from one genome to another (for example from Nipponbare *japonica* to an *indica* varietal group represented by IR 64). The workflow is as follows: (1) Get flanking sequences surrounding the target SNP in source genome (the main reference Nipponbare), (2) align these flanking sequences to target genome of variety of interest to verify if it hits a unique region in the target genome of similar location from the source genome, allowing some mismatches but not allowing multiple region hits, and (3) identify the flanking sequences surrounding the lifted-over SNP in the context of the target genome, for SNP assay design. The shared workflow is published in Rice Galaxy as [SNP lift-over].

#### 3k RG data access

Rice Galaxy provides access to the raw variant call format (VCF) files of each accession in the 3K RG project via connection (as data source in Rice Galaxy) to the 3,000 rice genomes at Amazon Web Services (AWS) Public Data (https://aws.amazon.com/public-datasets/3000-rice-genome/), with tools allowing region-specific download. In Rice Galaxy, tools in the [Get Data / FROM 3KRG] section allows listing of the accessions in the 3K RG and retrieval of genotype data for a selected accession of interest from the 3K RG collection. The subset genome region of interest (chromosome name – base start – base end) can be specified and extracted from the VCF of the accession of interest stored in AWS Public Datasets.

In addition, we developed an original Rice Galaxy component called Rapid Allelic Variant extractor (RAVE), which allows simultaneous extraction of genotyping data from several accessions of the internal 3K RG resource. It relies on the PLINK software [20], which efficiently builds a user-adjusted genotyping submatrix from a compressed PLINK binary biallelic genotype table (bed file + bim, fam files). Users can customize the genotyping dataset vertically by choosing a subpopulation (*indica*, *japonica*, *aromatic, aus, tropical, temperate*, etc.) or setting a list of varieties, and horizontally by restricting variations with a list of genomic regions, or a list of gene names. Additionally, users can filter the SNP positions by specifying thresholds for missing data or minor allele frequency (MAF). The extracted VCFs can be directly generated by Rice Galaxy, stored as output into the history pane of the Galaxy interface, and can be reformatted to Hapmap, a versatile file format for further analyses such as marker (SNP) design, GWAS analyses, or visualization in within a JBrowse [21] genome browser (Vcf2jbrowse component). External SNP datasets can also be imported into Rice Galaxy and merged with 3k accessions in order to compare and look at the closest genotypes using SNiPlay [22] workflow.

#### GWAS analysis using TASSEL

Using this feature, it is relatively easy to construct a genotyping matrix for a subset of accessions from the 3K RG and connect associated phenotypic information to perform GWAS analyses online, with outputs being decorated with various graphical enhancements. For the 3K RG accessions, the subset 1M GWAS and 440K Core SNPs that is usable for GWAS is already available as shared dataset in Rice Galaxy (Figure 2). Researchers working on the 3K RG panel can generate new phenotyping data from their respective experiments, upload the phenotype data into Rice Galaxy, and then perform GWAS using the TASSEL bioinformatics tool [23]. The GWAS Rice Galaxy workflow implementing TASSEL and Multi-Locus Mixed-Model package for association studies is shared from SNiPlay at Rice Galaxy (Figure 3).

**Figure 2.**
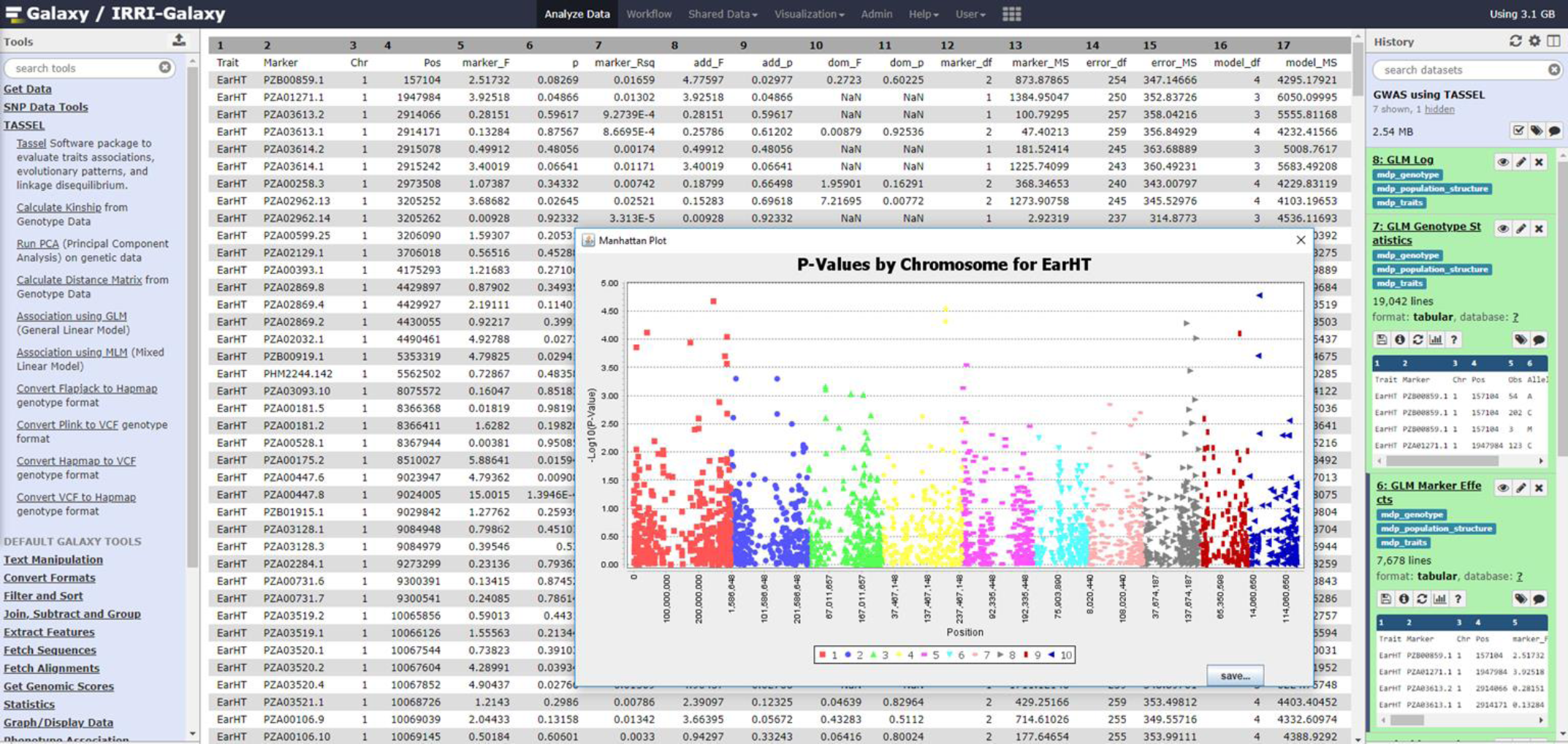
Genome-Wide Association Studies analysis (implemented by TASSEL software) in Rice Galaxy.

**Figure 3.**
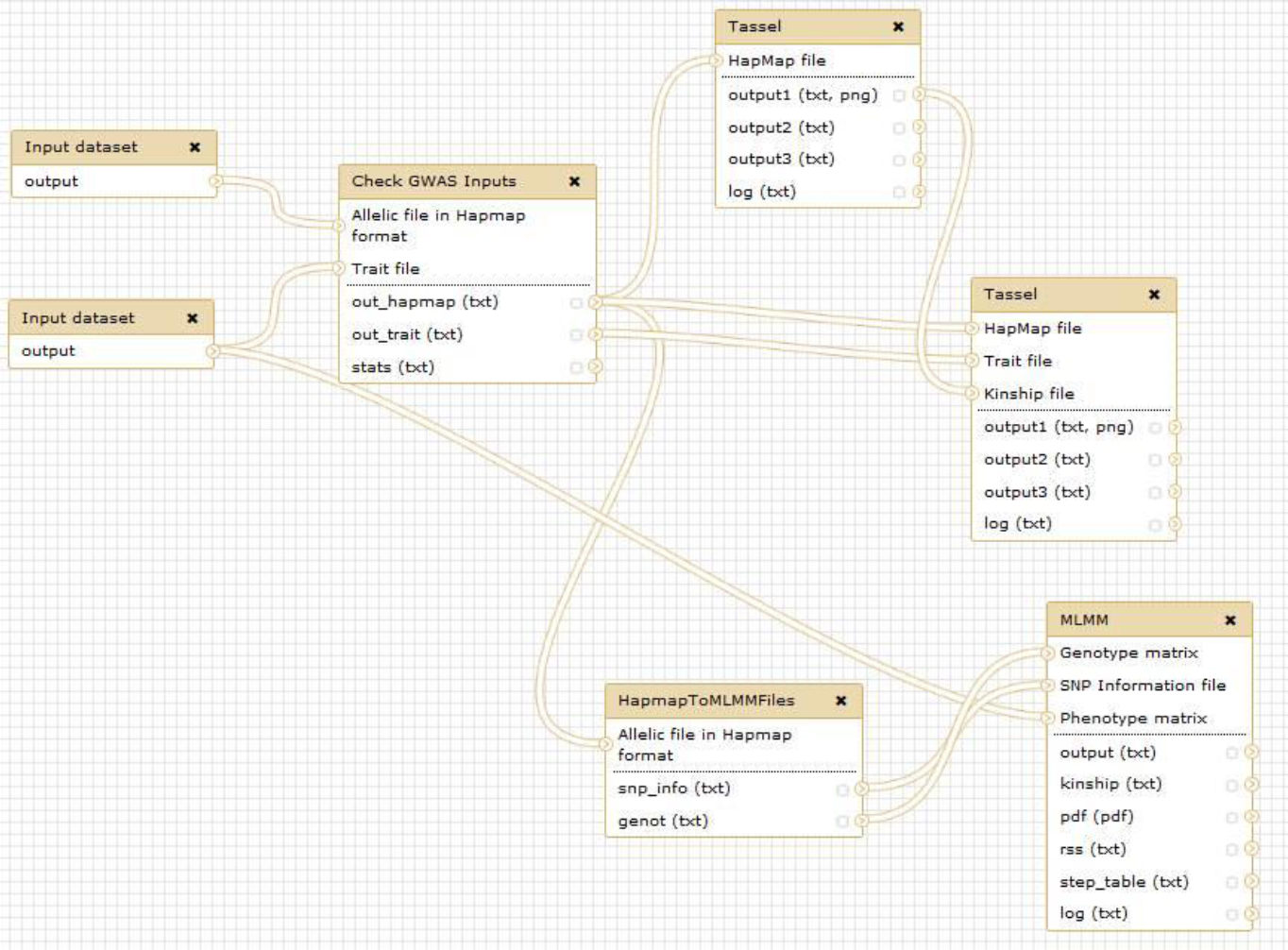
Genome-Wide Association Studies analysis workflow in SNiPlay as implemented in Rice Galaxy.

Aside from GWAS with 3K RG datasets, researcher-generated marker and phenotype data (outside of 3K RG) can also be uploaded to Rice Galaxy for GWAS analysis.

#### Genomic selection using Oghma genome prediction tool

Genomic selection (GS) is a promising breeding technique with potential to improve the efficiency and speed of the breeding process in rice [24]. With the intent of enabling the GS analysis process used on the 2 datasets in the Spindel et al. [24] study, (encoding data, filtering data to keep informative markers, creating a model from training set, evaluating the model and finally, performing the prediction itself), and to automate the analysis pipeline, the relevant packages (`methods, fpc, cluster, vegan, pheatmap, pROC, randomForest, miscTools, pRF, e1076, rrBLUP,` and `glmnet)` for the R Statistical language (https://www.r-project.org/) were installed in Rice Galaxy and the tool suite was collectively named Oghma (Operators for Genome decipHering by MAchine learning). Quality control tool (based on PLINK) and imputation tool using Beagle [25] (https://faculty.washington.edu/browning/beagle/beagle.html) were also installed. Four phenotype prediction/classifier methods (rrBLUP, random forest, SVM and lasso) were identified as relevant and deployed as tools in Rice Galaxy (Figure 4).

**Figure 4.**
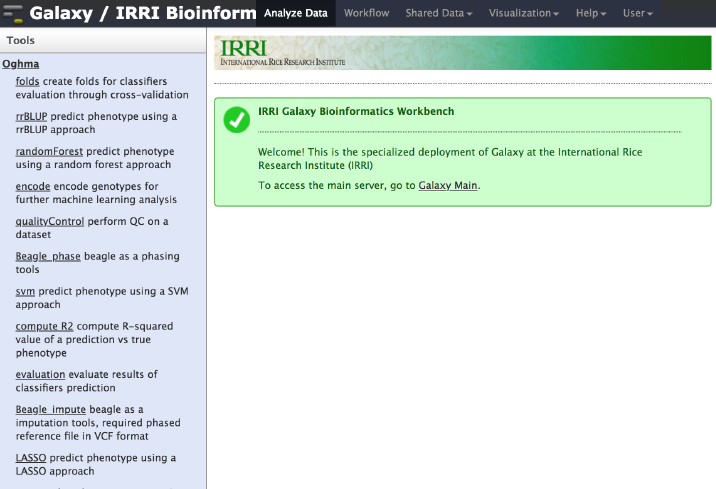
Oghma genomic prediction and selection tools in Rice Galaxy with various classifier tools installed.

Figure 5 shows the overall GS analysis workflow using Oghma. Genotypes are encoded through [encode] tool. For the training set, an encoded genotype and the corresponding phenotype files are used by a classifier tool to train a model, which can be used with another encoded genotype file to predict trait values (the genomic prediction). It is important to note that (1) both genotype for training and genotype to predict must have the same markers (and thus, genotype files must have the same number of columns) to make a prediction, and (2) the “evaluation” option of the classifier tool can have any value except 1 (it is recommended to keep the default value = 0).

**Figure 5:**
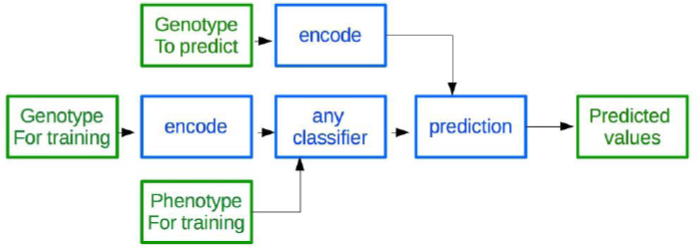
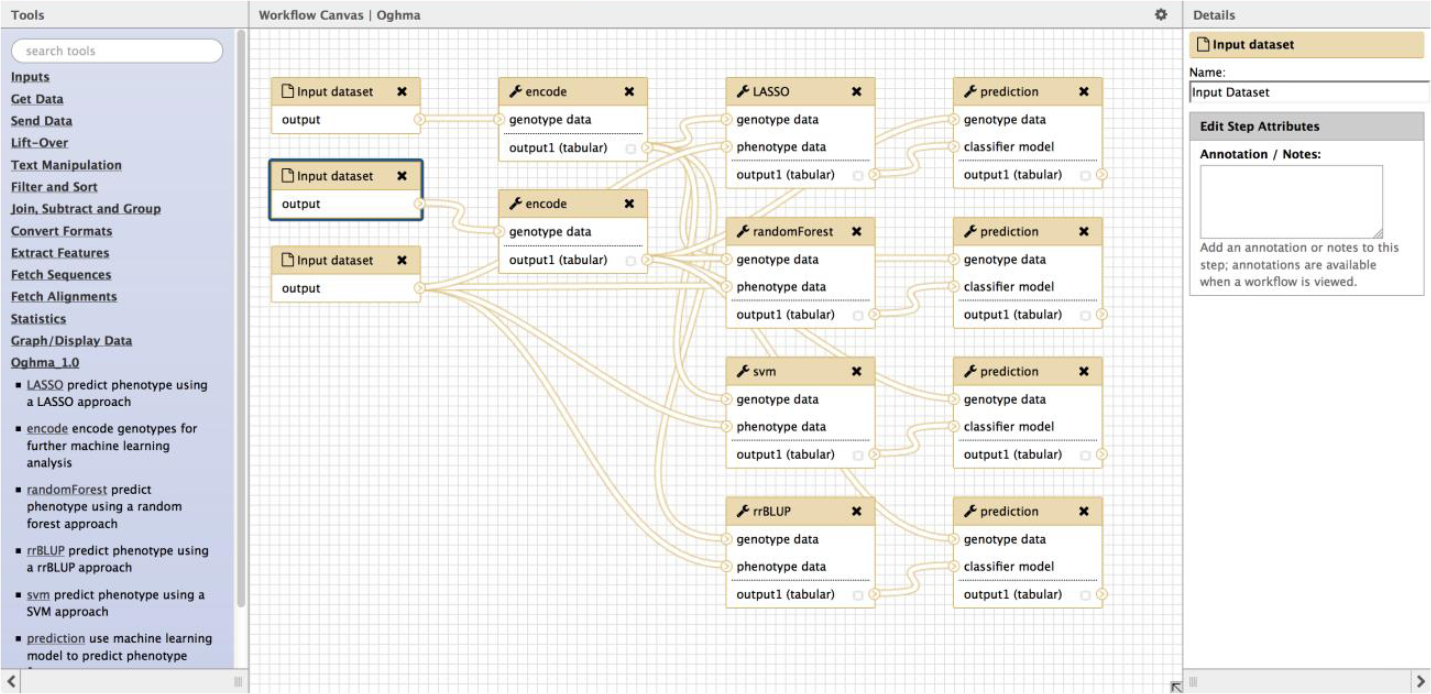
Genomic Selection analyses workflow as implemented by Oghma tool suite. A. Overview of the Genomic Selection analyses workflow as implemented in Oghma tool suite. B. Rice Galaxy workflow for genome prediction using Oghma tool suite.

A big challenge when using machine learning approaches for genomic prediction is the optimization of the model based on training data, specifically setting the best parameters of the methods mentioned prior. Oghma was designed to automate the optimization of the parameter(s) of the classifiers on the fly (as opposed to manual tweaking), thus allowing users without experience of machine learning to easily optimize a model for their own data. Oghma includes some tools to evaluate prediction accuracy to allow the user to choose the most accurate method on their data by performing a cross-validation with a user-uploaded training set. Two metrics, the coefficient of determination (R^2) and the correlation between predicted and observed phenotype, and a visualization (scatterplot of predicted vs observed) have been implemented to evaluate the methods. The [computeR2] and [plotPrediction] tools are used to compute R^2 and visualize the accuracy of prediction. These tools both take the true phenotypes and the predicted outputs as inputs (take note that both predictions and phenotypes data must be in the same order), and return the computed R^2 or the scatterplot display of true phenotype vs prediction. Oghma can be used to evaluate a classifier (Figure 6). Like the general GS workflow, genotype and phenotype are used as input for any classifier, but the “evaluation” option must be set to 1. Fold for cross-validations are designed through the [fold] tool, which take as input the encoded file. These folds are used as extra argument by the classifier tools. The chosen classifier tool produces a file, which is not a model but the prediction of the test set for each cross-validation. This output is used as input, along with the phenotypes and folds, by [evaluation], which output some performances index (R^2 and correlation). Although it does give a real indication of performances, trying to predict the training set (i.e. using the same genotype file in the pipeline described above), or at the least, showing if the classifier is not under-fitting the data.

**Figure 6.**
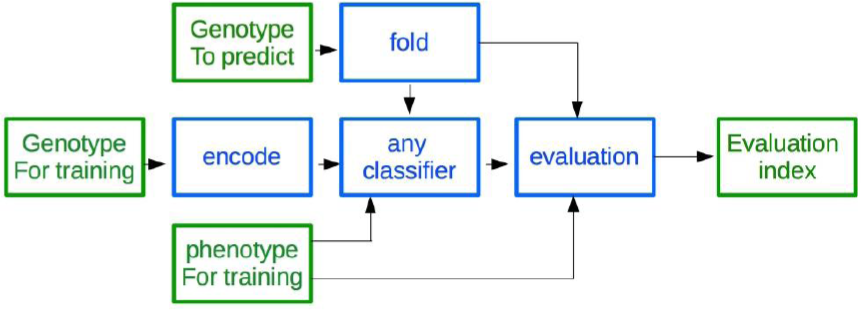
Workflow for classifier evaluation in the genome prediction tool suite implemented by Oghma.

We installed several classifiers in Oghma to allow users to test the best one(s) suited for their dataset, as our literature survey shows that no method seems to outperform the others on all genomic prediction tasks. It was noticed that Random Forest was the most accurate and the most stable classifier on Spindel dataset [24], thus we set this as default in Oghma. An original aggregation method is also implemented in Oghma, aggregating outputs of multiple classifiers to improve prediction. This tool takes as input the prediction of *n* classifiers and tries to aggregate them through weighted mean of the prediction (weight optimized by genetic algorithm) or regression (multiple type of regression have been implemented, based on decision tree, SVM and Random Forest). Limited testing shows that this approach is promising, matching Random Forest in some cases, especially with a meta-SVM, with polynomial or linear model, as aggregation method, but still needs some improvement as the accuracy remains unstable when evaluated through cross-validation (data not shown). The aggregation method can also be evaluated using the aforementioned evaluation tools.

#### Diversity and population structure analysis of end-user datasets

SNP datasets – such as those extracted from the 3K RG resource after a filtering by the RAVE module or custom sets directly uploaded in Rice Galaxy environment (Figure 7) can be processed for a complete exploration and large scale analysis thanks to the SNiPlay Rice Galaxy workflow (Figure 8). The workflow is available through the instance, requiring a VCF file as input. This workflow allows various analyses: (i) SNP annotation by snpEff (http://snpeff.sourceforge.net/) wrapper preconfigured for RGAP release 7.0 [26] (http://rice.plantbiology.msu.edu/) gene models (ii) variant filtration using PLINK wrapper, (iii) general statistics such as Transition-Transversion ratio, levels of heterozygosity and missing data for each variety using VCFtools, (iv) SNP density analysis, (v) diversity indices calculation in sliding windows along the genome using VCFtools (Pi, Tajima’s D, FST if subpopulations provided), (vi) linkage disequilibrium, (vii) population structure by sNMF (http://membrestimc.imag.fr/Olivier.Francois/snmf/index.htm), (viii) Principal Component Analysis and Identity By State (IBS) clustering of varieties by PLINK, and (ix) SNP-based distance phylogenetic tree by FastME (http://www.atgc-montpellier.fr/fastme/). Most key steps are decorated with sophisticated visualizations using a dedicated plugin. Visualization can be displayed by clicking on the [visualization] icon.

**Figure 7.**
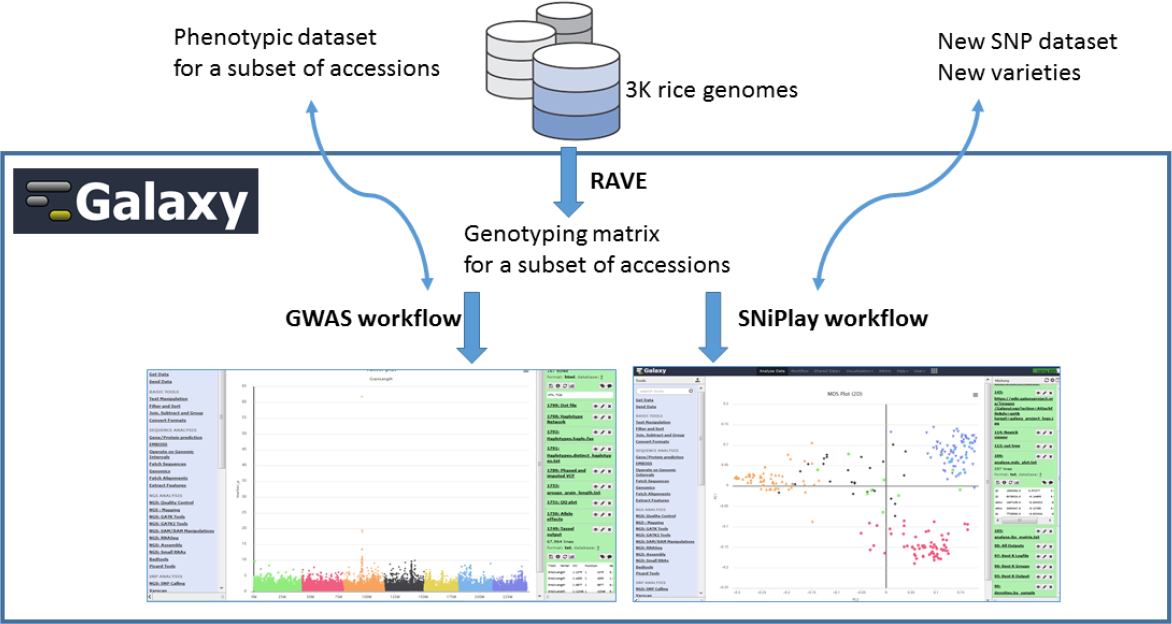
Overview schematic showing the integration of the 3K Rice Genomes project genotyping database and rapid extraction of subset SNPs by RAVE module for use by analyses workflows installed in Rice Galaxy.

**Figure 8.**
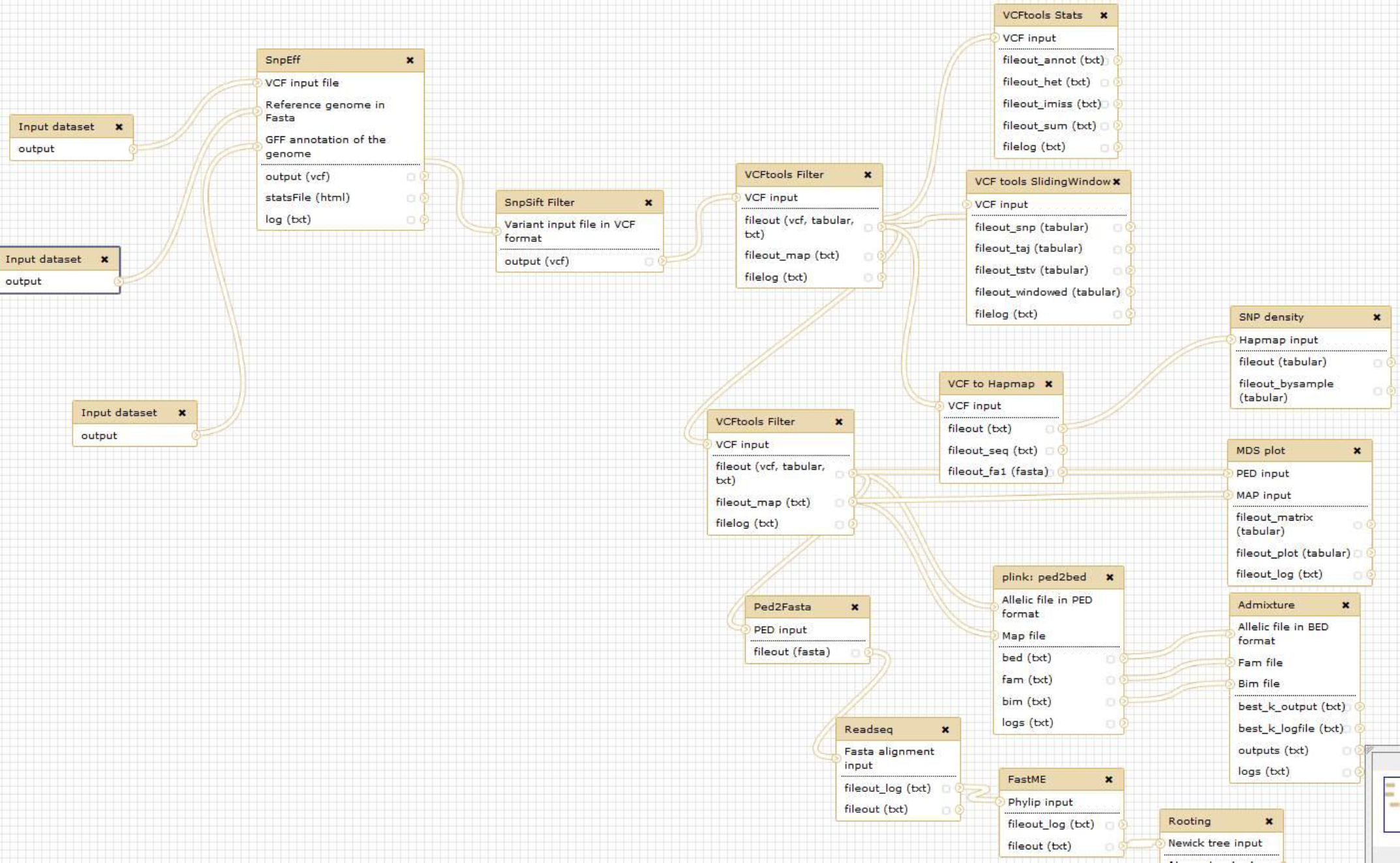
Rice Galaxy SNiPlay workflow for diversity and population structure analyses using various software tools.

In practice, this workflow can be processed for many applications such as the identification of possible introgression events, the identification of putative genomic regions involved in the control of qualitative trait through a FST approach, the investigation for potential duplicates in the 3K RG accessions dataset and custom datasets, or the estimate of closest varieties of new sequenced accessions, by ranking a list of varieties from the database most closely matching the given sample. It can be used also for the close inspection of genomic region of interest after a GWAS analysis, through a linkage disequilibrium focus or the haplotyping of candidate genes.

#### Uniqprimer module

Uniqprimer is a workflow for comparative genomics-based diagnostic primer design, developed from a pipeline used in-house at Colorado State University to develop novel species and subspecies-level diagnostic tools for bacterial plant pathogens including pathovars of *Xanthomonas translucens* [27], geographical variants of rice-associated *Xanthomonas* spp. [28–30], and the genetically diverse rice pathogen *Pseudomonas fuscovaginae* [31]. Uniqprimer is now deployed in Rice Galaxy for user-friendly diagnostic primer design from draft or complete pathogen genomes. The user inputs multiple bacterial genomes from diagnostic target species as well as non-target species (i.e. “include” and “exclude” genome files), and the tool performs comparative alignment, primer design, and primer validation to output a list of primers that are specific to the target genomes (Figure 9). The uniqprimer standalone program is written in Python and is available at the Southgreen github repository (https://github.com/SouthGreenPlatform/Uniqprimer), along with the detailed documentation for developers and end-users.

**Figure 9.**
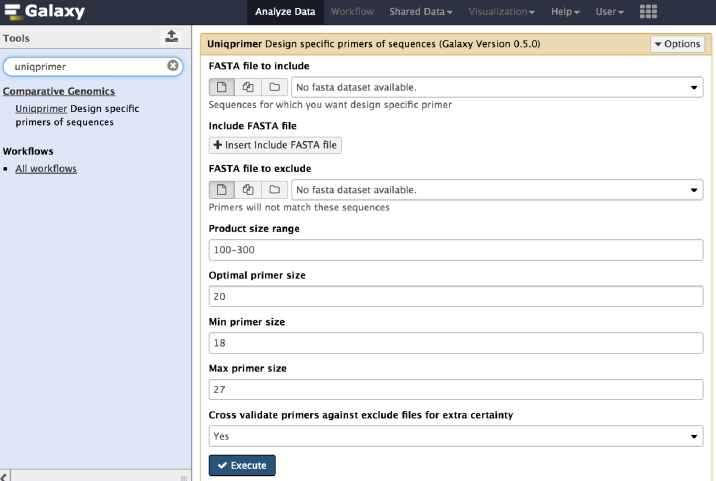
Uniqprimer comparative genomics-based diagnostic primer design tool for microbial pathogen detection installed in Rice Galaxy.

### 3. Rice Galaxy OA: a Prototype for Open Access

IRRI, as a member center of the Consultative Group for International Agricultural Research (CGIAR, https://www.cgiar.org/), complies with the CGIAR policy on Open Access and Open Data (https://www.cgiar.org/how-we-work/accountability/open-access/). In collaboration with Indiana University in the United States and National Institute of Advanced Industrial Science and Technology in Japan, and carried out through grants from the National Science Foundation (NSF) in the US and the MacArthur Foundation through the Research Data Alliance (RDA – https://www.rd-alliance.org/), the team undertook a prototyping effort to bring the Rice Galaxy system to maximum compliance with the CGIAR policy.

The basis for the design to add open access to Rice Galaxy is a foundational technical idea emerging from activities occurring in the international RDA. This idea acknowledges that for open data access to be broadly realized, all meaningful data objects must have a globally unique and persistent identifier (PID). Globally unique means the name is not shared with other objects on a global scale. An identifier is persistent when the PID itself cannot be destroyed, and when the relationship between the identifier and the data object it points to is permanent. Through an international working group in RDA, a team of researchers is advancing the notion of PID Kernel Information, which injects a tiny amount of carefully selected metadata into a PID record. This technique has the potential to stimulate an entirely new ecosystem of third party services that can process the billions of expected PIDs. The key challenge of this working group is to determine which from amongst thousands of relevant metadata are suitable to embed in the PID record.

Our design draws on earlier work by us in data provenance capture and representation [32–34] and employs a hands-off technique (*data provenance capture*) to gather information about a researcher’s rice genomics analysis as the analysis is running. Through this technique, information acquired while the analysis is running is compiled and combined with pre-analysis information that is available at the beginning of the analysis workflow. Such information includes who performed the analysis, when it was performed, and under what conditions.

There have been earlier approaches to capture provenance of Galaxy workflows. Geocks *et.al* [35] developed a history panel for users to facilitate reproducibility. Gaignard *et al*. [36] proposes the SHARP toolset, a semantic web (i.e. linked data) approach of harmonizing provenance collected from both the Galaxy and Taverna workflow systems. Kanwal *et al*. [37] captured the activity of a workflow (called a *provenance trace*) including the version of analysis tools run, the software parameters used, and the data objects produced at each workflow step. This work also targets increased reproducibility of past workflow instances. Missier *et al.* [38] proposes the “Golden Trail” architecture to describe and store workflow runtime provenance retrieved from Galaxy. The golden trail of provenance that is collected can be used to construct a virtual experiment view of past workflow runs. The four research contributions described further underline the need for the capture of provenance from workflow systems. They propose different but equally important uses of data provenance, that is, to facilitate the improvement of science through reproducibility and construction of virtual views of an experiment once it has completed.

Our design for Rice Galaxy Open Access (OA) shares similarities with these other techniques, however its end goal is different, which is to advance open access, hence making Rice Galaxy consistent with CGIAR’s open access policy. To do this, we focus on each piece of data and information deemed valuable that emerges from workflow runs deemed to be of importance. This particular data and information must be retained and shared with others, while being subject to reasonable restrictions. This is a highly selective approach to provenance capture, and one that makes our work unique. We briefly outline the solution here and identify resources for those interested in pursuing the topic in more detail.

The architecture of Rice Galaxy OA (Figure 10A) utilizes the Handle system [39] and two standards emerging from the Research Data Alliance, RDA PID Type [40] and the Data Type Registry [41]. It additionally uses storage and compute resources provisioned through the NSF funded project, Pacific Rim Applications and Grid Middleware Assembly (PRAGMA).

**Figure 10.**
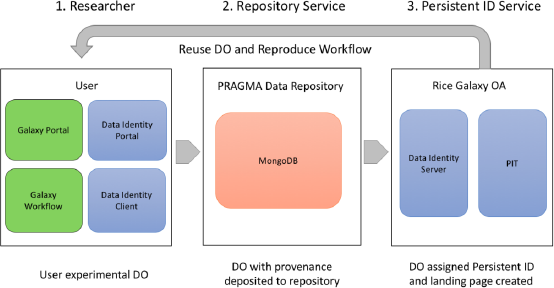
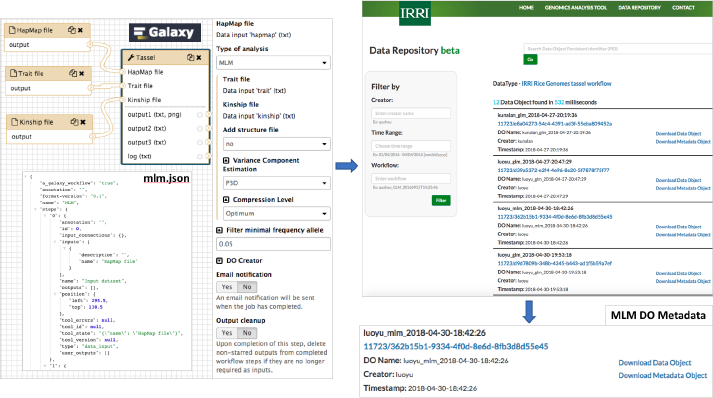
The components (A) and the flow of Digital Objects from upload to discoverability (B) in the prototype Rice Galaxy Open Access. A. The underlying software infrastructure for the components of Rice Galaxy Open Access. B. Digital Object flow in Rice Galaxy Open Access. A Galaxy analysis workflow (exported as JSON file) is deposited to the DO repository, and the data identity server publishes the deposited DO + meta-data for discoverability.

A researcher interacts with the open access enhanced Rice Galaxy system as follows:

1. Researcher performs an analysis in Rice Galaxy
2. Data objects (input data, output data, information such as configuration parameters) are extracted from Rice Galaxy OA into a PRAGMA Data Repository Database (MongoDB) (Figure 10A),
3. The data objects are assigned Persistent Identifiers, the PID Kernel Information is assigned into the PID record at this time, and a landing page created for each (Figure 10B).
4. Data objects can be downloaded from the Data Identity server and re-loaded to the Rice Galaxy server for full faithful reproduction of the analysis

The resulting system appears to be promising and addresses a number of the recommendations from CGIAR. The Rice Galaxy OA system is a user transparent means of harvesting digital objects from applications and assigning PIDs to scientific outcomes. The architecture is modular and built with default PID information types and metadata using RDA products (Figure 10A). Although this proof-of-concept prototype successfully demonstrates the feasibility of this approach, there remains some future work. The community needs to provide feedback on which data and information products are most important to retain and make available. Additionally, not all workflow runs are important to a researcher as they could be system tests or new workflow tests. Thus, how a researcher identifies the items he/she wishes to make available to others and when, remains an important consideration for this system. For more information, points of contact to the team, the underlying software for Rice Galaxy OA, and the link to the prototype server can be found at https://github.com/Data-to-Insight-Center/RDA-PRAGMA-Data-Service/wiki/Welcome-to-PRAGMA-Data-Service-Prototype.

### 4. Rice Galaxy architecture discussion

We deployed IRRI Galaxy in an AWS EC2 instance (t2.large instance 2 vCPU, 4 GiB RAM) for the production server deployment in the cloud with Linux Ubuntu release 12.04.2 LTS (GNU/Linux 3.2.0-40-virtual x86_64) using Galaxy release 14 as described in the Galaxy documentation.

External data from the 3K RG Project files stored in the 3K RG AWS Simple Storage Service (S3) Public Data resource hosted at http://s3.amazonaws.com/3kricegenome/ (or s3:// 3kricegenome/) is accessed using AWS S3 Command Line Interface, a command line tool utility in AWS that provides an interface to access AWS S3 objects (CLI, https://docs.aws.amazon.com/cli/latest/reference/s3/). First, Rice Galaxy connects to the 3K RG AWS bucket using s3API and allows the objects inside the bucket to be transparent to Galaxy. VCF files (and the accompanying index files) are downloaded to Rice Galaxy using the S3 CLI with the `aws S3 cp` command, executed as:

~~~
aws -profile user s3
cp s3://3kricegenome/REFERENCE/VCF_FILE.snp.vcf.gz*.
~~~

The subset region of the VCF file (chromosome:start-end) is then extracted using BCFtools (http://samtools.github.io/bcftools/) wrapped in Rice Galaxy and exported to the history pane as bgzipped, indexed BCF file, which can then be converted back to VCF using [VCFTOOLS] in Rice Galaxy. Standard methods for tool wrapper development and deployment were followed. All tool wrapper XMLs developed specifically for Rice Galaxy are deposited and shared in a project-specific Rice Galaxy toolshed repository at http://52.76.88.51:8081/ (Figure 11) and will also be deposited in the central Galaxy toolshed (https://toolshed.g2.bx.psu.edu/). All developments and testing of Rice Galaxy and Rice Galaxy Toolshed were done in Docker containers hosted in virtual machines at the Advanced Science and Technology Institute, Department of Science and Technology of the Philippine Government (ASTI - DOST) prior to final deployment to the AWS instance.

**Figure 11.**
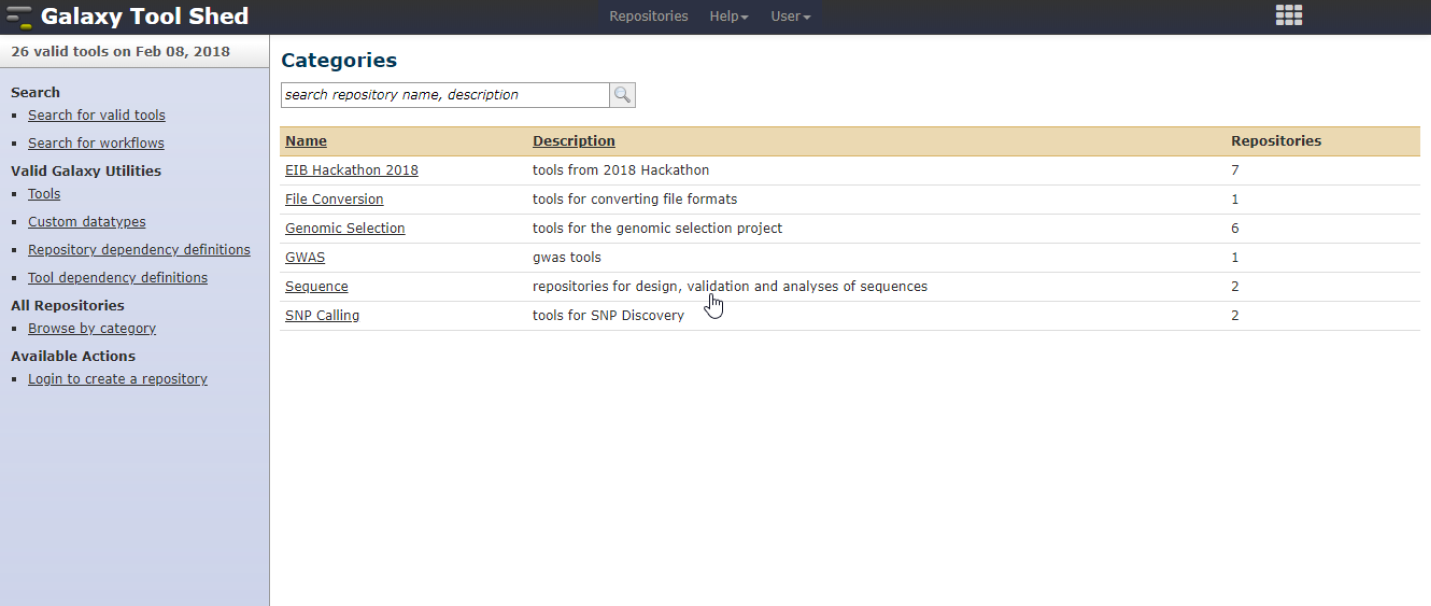
Rice Galaxy Toolshed with the various available tools.

This resource will empower the rice research community to benefit from publicly available datasets (e.g. 3K RG) and materials (seed/accessions), to enhance or even drive their own respective institutional genetic/genomic/breeding efforts. The Rice Galaxy instance (data, tools, computing resources) is free for use by all.

In addition to the integration of these tools, new Galaxy wrappers and visualization plugins are being developed for visualizing chromosomes and their information (SNP density, structural variants, translocations) either in linear or circular mode, using recent web technologies (Ideogram.js [42], BioCircos.js [43], respectively).

Finally, a Docker container of Rice Galaxy is under development so that it can be easily shared and deployed through the Galaxy Docker flavor initiative (https://github.com/bgruening/docker-galaxy-stable).

## Conclusion

Rice Galaxy is a federated Galaxy resource specialized for rice genetics, genomics, and breeding. The resource empowers the rice research community to utilize publicly available datasets (3K RG), materials (seed/accessions), and their own data, allowing complex data analyses to be performed even without investment in their own computational infrastructure and software development team. Rice research – related tools are also hosted in Rice Galaxy (i.e. Uniqprimer rice pathogen diagnostic design).

Rice Galaxy is freely accessible to all and we invite the rice research community to participate in enriching the tools hosted by the resource. It can serve as a repository for data, analyses results, and new bioinformatics tools coming from institutions that have used the publicly available rice diversity panels from 3K RG, or have developed rice genomic/genetic analyses tools that they wish to share to the community, and a computing infrastructure for small institutes without in-house computing capability.

### Availability and requirements

Project name: RICE GALAXY

Project home page: https://github.com/InternationalRiceResearchInstitute/RiceGalaxy

Operating system(s): Linux Ubuntu release 12.04.2 LTS

Programming language: Python

Other requirements: R release 3.2.3 and following packages: methods, fpc, cluster, vegan, pheatmap, pROC, randomForest, miscTools, pRF, e1076, rrBLUP, glmnet ;TASSEL release 5.2.40; plink v1.90b3k; JBrowse 1.14.1; snpEff 4.3T; sNMF 1.2 (and as R package LEA); FastME 2.0

License: Rice Galaxy tools are released under GNU GPL. All software from external sources is bound by their respective licenses.

Any restrictions to use by non-academics: Rice Galaxy tools are without restriction to non-academics. All software from external sources is bound by their respective non-academic restrictions

Code availability: Tool wrappers at Rice Galaxy Toolshed (http://52.76.88.51:8081/). Rice Galaxy is available at IRRI Github (https://github.com/InternationalRiceResearchInstitute/RiceGalaxy).

### Availability of supporting data

3,000 Rice Genomes Project at Gigascience database (http://gigadb.org/dataset/200001)

3K RG BAM and VCF files available from Amazon Public data and ASTI-DOST IRODs site, instructions at http://iric.irri.org/resources/3000-genomes-project.

SNP sets and morpho-agronomic characterization from 3K RG at SNP-Seek download site (http://snp-seek.irri.org/_download.zul)

### Availability of supporting source code and requirements

Project name: Uniqprimer

Project home page: https://github.com/SouthGreenPlatform/Uniqprimer

Operating system(s): Linux OS

Programming Language: Python

Other requirements: MUMmer 3

License: GNU GPL

Project name: PRAGMA Data Service

Project home page: repository https://github.com/Data-to-Insight-Center/RDA-PRAGMA-Data-Service/wiki/Welcome-to-PRAGMA-Data-Service-Prototype

Operating system(s): Platform independent

License: Apache License 2.0

## Declarations

### Abbreviations

3K RG: 3,000 Rice Genomes
HDRA: High Density Rice Array
SNP: single nucleotide polymorphism
GWAS: Genome-Wide Association Studies
RAM: random access memory
CPU: central processing unit
IRRI: International Rice Research Institute
NGS: next-generation sequencing
QTL: quantitative trait loci
IRGSP: International Rice Genome Sequencing Project
RGAP: Rice Genome Annotation Project
KASP: Kompetitive Allele Specific PCR
VCF: variant call format
AWS: Amazon Web Services
RAVE: Rapid Allelic Variant extractor
MAF: minor allele frequency
TASSEL: Trait Analysis by aSSociation, Evolution and Linkage
GS: genomic selection
Oghma: Operators for Genome decipHering by MAchine learning
rrBLUP: ridge regression best linear unbiased predictor
SVM: support vector machine
FST: fixation index
NSF: National Science Foundation
CGIAR: Consultative Group for International Agricultural Research
RDA: Research Data Alliance
PID: persistent identifier
OA: open access
PRAGMA: Pacific Rim Applications and Grid Middleware Assembly
EC2: elastic computing cloud
CLI: command line interface
S3: Simple Storage Service
API: Python Application Programming Interfaces
XML: eXtensible Markup Language

### Competing interests

The author(s) declare that they have no competing interests.

### Funding

Components of the project are supported by the following grants: Taiwan Council of Agriculture Grant to IRRI, International Rice Informatics Consortium, and CGIAR Excellence in Breeding Platform for financial support to the Rice Galaxy main server, the USA National Science Foundation PRAGMA grant number: NSF OCI 1234983, the RDA/US-sponsored adoption program funded by the MacArthur Foundation, and the AIST ICT International Collaboration Grant.

### Authors’ contributions

VJ and AD equally contributed to create Rice Galaxy. NB contributed the genomic prediction tools. AD, GD, PL, and MR contributed the RAVE and SNiPLAY tools. JD, JRM, JPP created the development Rice Galaxy cloud instances hosted at DOST-ASTI. LM created the SNP-Seek interfaces. LT, JL, JEL contributed the Uniqprimer tool. GZ, KR, BP, and JH contributed the Rice Galaxy Open Access, MT, NA, and TK contributed to funding acquisition and writing, RM coordinated the conceptualization of the project and the writing process.

#### Acknowledgments

The authors are grateful to the following people and institutions/agencies for their support: DOST-ASTI for hosting the Rice Galaxy toolshed server, Jay Santos and Denis Diaz for assistance with AWS architecture.

